# How partial phenotyping to reduce generation intervals can help to increase annual genetic gain in selected honeybee populations

**DOI:** 10.1101/2024.10.30.621079

**Authors:** Tristan Kistler, Evert W. Brascamp, Benjamin Basso, Florence Phocas, Piter Bijma

## Abstract

Honeybee breeding is organized around annual cycles, following seasonal change. Generation intervals are thus commonly multiples of whole years. Most queens are generally raised during spring or early summer in temperate climates. A generation interval of 1 year limits phenotyping to early recordable traits, before the spring following queens’ births. Some traits, however, can only be recorded later, as is typically the case for total honey yield. Their recording on selection candidates thus increases the generation interval to at least 2 years, a common interval on the dam path. Using stochastic simulation, we investigated the impact of halving the dam generation interval and therefore recording only early traits on candidate dams. The generation interval on the sire path remained at 2 years with complete phenotyping. Breeding goals with varying weights on early and late traits were considered, as well as different genetic correlations between traits, from negative to positive. The acceleration of the breeding scheme generally resulted in an increased genetic gain for the two-trait breeding goals, from 0% up to +47% after 20 years of selection. Although inbreeding rates per generation were slightly lower in the accelerated breeding scheme, associate inbreeding levels were about 20-30% higher after 20 years of selection due to the 33% increased generation turnover. To avoid too high inbreeding, shortening the generation interval should thus be accompanied by strategies to limit inbreeding while still retaining most of the genetic gain, such as increasing the breeding nucleus size by relaxing selection intensity.

## Introduction

Beekeeping is facing major challenges, with global threats such as climate change, pesticides, and the spread of parasites and diseases, driving high colony losses (Steinhauer *et al*., 2018; Hristov *et al*., 2020; Insolia *et al*., 2022). As a result, colony replacement and artificial queen rearing has become more intensive to replace lost or unproductive stock (Aureille, 2014; Kouchner *et al*., 2019). Possibly having to actively choose breeding queens instead of letting colonies spontaneously requeen has created awareness for selective breeding among beekeepers. Selection programs are thus emerging worldwide with a variety of breeding schemes. In these breeding schemes, bee breeders generally aim to improve a collection of traits simultaneously, focusing more or less on colony production, manageability, and resilience (Guichard *et al*., 2019; Maucourt *et al*., 2020; Hoppe *et al*., 2020; Kistler *et al*., 2024). They generally aim to obtain enough genetic gains in both the short and long term while minimizing inbreeding rates to ensure the sustainability of their breeding programs.

Honeybee breeding is usually organized around annual cycles, following seasonal change. Possible generation intervals are thus commonly of 1-year increments, typically from 1 to 4 years (Uzunov *et al*., 2022; Basso *et al*., 2024). Most queens are generally raised during spring or early summer in temperate climates. This is mainly because queens mate shortly after birth, and drones (males) are most reliably produced by colonies in that period (Page, 1981). They then store semen in their spermatheca and will not mate anymore after the onset of egg-lay, a few days or weeks later. Performance testing can only start months later, once queens have established new colonies (Büchler *et al*., 2024). A generation interval of 1 year thus limits phenotyping to early recordable traits, before the spring following queens’ births. These early traits might include hygienic behavior (Büchler *et al*., 2020) linked to brood diseases; manageability traits (gentleness and calmness); first winter survival, winter brood interruption, winter feed consumption or early spring colony development; and possibly partial honey yield depending on floral resources. Complete honey yield, however, can generally only be obtained after the following mating period, when queens are over 1 year old. Other important traits, such as swarming tendency, can show significant variability only in colonies with older queens (Uzunov *et al*., 2014). To allow for complete colony phenotyping of candidates, most breeding schemes thus use a 2-year generation interval in dams. However, some use partial phenotyping to reduce it to 1 year (Basso *et al*., 2024; Kistler *et al*., 2024).

Due to the high mortality rates in honeybees, reduced generation intervals can also increase selection intensity and genetic diversity by selecting queens in a bigger candidate population before their second winter. In addition, shortening generation intervals is a well-known and powerful way to increase the expected annual genetic gain, which is inversely proportional to the generation interval when ignoring inbreeding and for a fixed selection accuracy (Dickerson and Hazel, 1944).

Predicting genetic gain and inbreeding is challenging using deterministic derivations, especially when considering the haplo-diploidy and polyandry in honeybees, as well as the complex colony phenotypes that result from two genetic effects expressed by two different generations (the queen and its worker group). Comparing breeding strategies can more accurately be done by stochastic simulations, considering bee specificities in addition to detailed representations of breeding populations, including limited population sizes, overlapping generations, and unbalanced sizes of sister groups due to random mortality. In addition, the effect of limited genetic connection across contemporary groups (apiaries and years) on genetic parameters and breeding values estimation can also be integrated (Clément *et al*., 2001). Stochastic simulations are therefore useful to assess both expectations and standard deviations of the genetic gain and inbreeding level in bee breeding schemes.

Using stochastic simulation integrating bee specificities, Du et al. (2023) studied the shortening of the sire’s generation interval from 3 to 2 years in a single-trait simulation study. Assuming genetic parameters known, they showed that the reduction in the generation interval brought more genetic gain (about +12% after 20 years) but also a more than 2-fold increase in mean inbreeding levels and rates per year. However, the reduction of the generation interval was studied with selection for a single trait that was recorded for all alternative schemes. Furthermore, their assumption of known genetic parameters is likely to have underestimated the impact of parameter estimation on the variability of results (Du *et al*., 2022a, 2022b). Estimation of genetic parameters is particularly challenging in honeybees as queen and worker effects are statistically highly confounded and populations usually have breeding nuclei of only a few tens of breeding queens (Maucourt *et al*., 2020; Guichard *et al*., 2020; Basso *et al*., 2024; Kistler *et al*., 2024).

In this study, we will assess by stochastic simulation the impact of reducing the dam’s generation interval from 2 to 1 year in a two-trait selection program involving an early and a late recorded trait. The reduction in the dam generation interval meant that only the early trait could be phenotyped for candidate dams, while for candidate sires, complete phenotyping was maintained with a two-year generation interval. Results in terms of genetic gain and inbreeding will be contrasted with a breeding scheme maintaining complete phenotyping and a 2-year generation interval on both dam and sire paths. Various breeding goals and genetic correlations between the early and late traits, as well as between worker and queen effects, are covered.

## Methods

### Genetic model

Simulations implemented an infinitesimal genetic model with individual queens, drones, and worker groups, with two additive genetic effects affecting colony phenotypes (y): queen genetic effect expressed by the colony’s queen 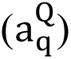 and the average worker genetic effect expressed by the colony’s worker group 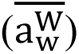 (Kistler *et al*., 2021):

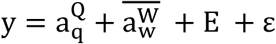

where E is an environmental apiary by year effect and ‘ε’ is a random residual term. Simulation of apiary effects is described further below.

Two traits were simulated: an early and a late-recordable trait. The main equations for the generation of base individuals’ breeding values (BVs) and of their inheritance are given in File S1.

### Breeding goals

Breeding goals (H) were defined as weighted sums of true BVs for each trait, using 5 combinations of weights on both traits. The weight combinations varied from placing all the emphasis on the early trait to all the emphasis on the late trait in 25% increments. The resulting combinations were: (1:0), (0.75:0.25), (0.5:0.5), (0.25:0.75), and (0:1), where the first number is the weight put on the early trait and the second on the late trait. The weight applied to each trait was added as a subscript to H (e.g., H_0.25:0.75_). The BVs in H for worker and queen effects were summed together with the same weight. The selection index combined the estimated breeding values (EBVs) of each trait (or phenotypes in the Initialization phase, see further below) with the same weights as the corresponding breeding goal. EBVs for worker and queen effects on the same trait were summed before applying the weights in the selection index as in the breeding goals. Dams used to produce queens (from fertilized eggs) were selected based on their average offspring EBVs (i.e. the EBVs of their worker group). Sires used to produce drones (from unfertilized eggs) were selected based on their own EBVs (Brascamp and Bijma, 2014; Brascamp *et al*., 2016).

### Simulation process

Each simulation replicate was divided into three phases: an ‘Initialization’ phase, followed by a ‘Base breeding scheme’ (Base), and an ‘Alternative breeding scheme’ (Alt), both building upon the Initialization outcomes (Fig. 1). The Initialization phase generated a closed population under selection and accumulated pedigrees and phenotypes for the subsequent phases. Base followed the same structure as Initialization with a generation interval of 2 years on both the dam and the sire path. In contrast, in Alt, the dam’s generation interval was reduced from 2 to 1 year, limiting phenotyping to the early trait only and resulting in an average generation interval of 1.5 years (25% less than in Base). The three phases will be detailed next.

**Fig. 1.**
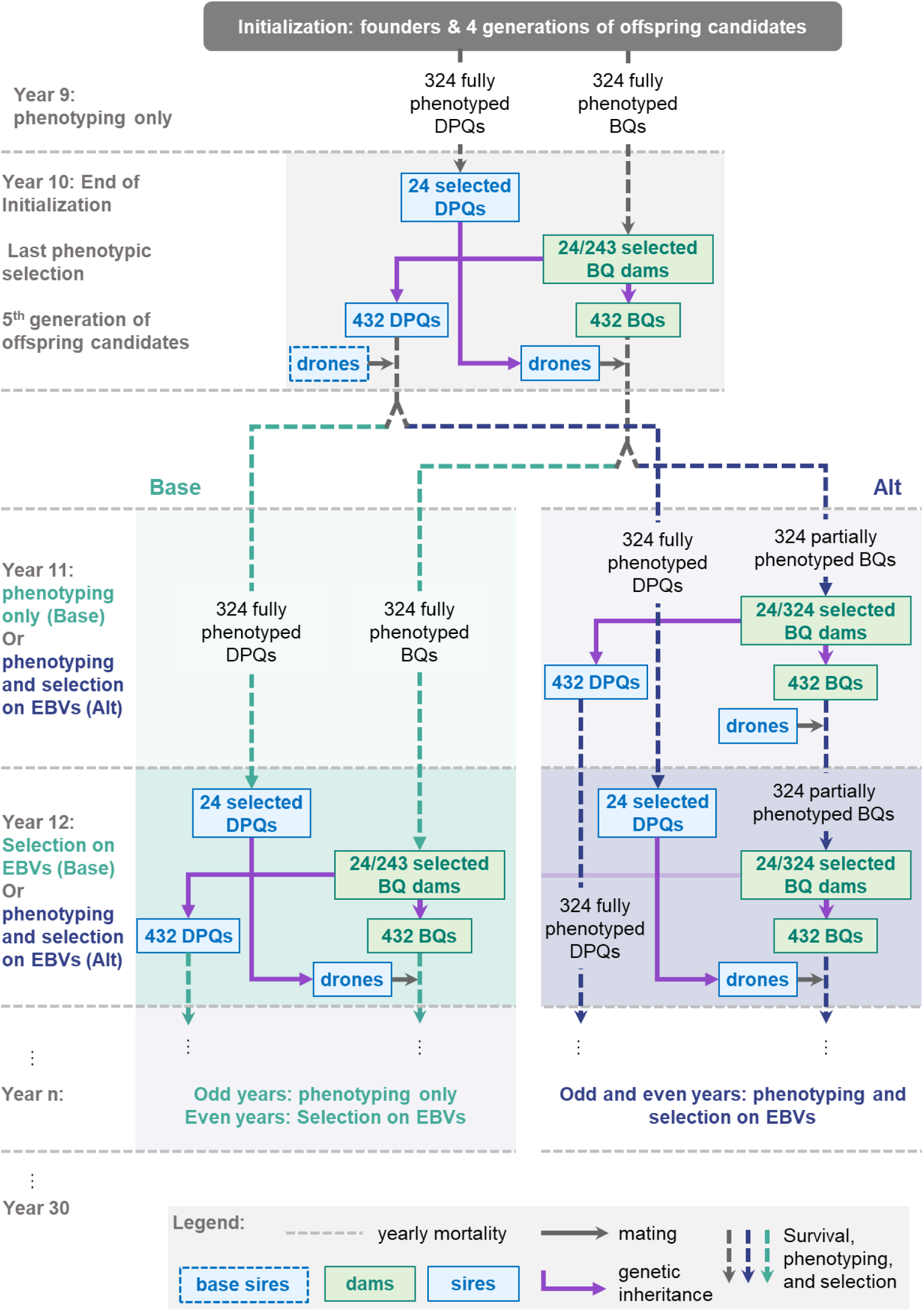
Simulation process: Initialization, followed by the Base and Alternative breeding schemes. BQ: breeding queen. DPQ: drone-producing queen. BQ refers to queens that were candidates for selection, and “selected BQ” to the selected dams. See main text for a description of the simulation process.

#### Initialization phase

In the Initialization, potential dams (BQs, breeding queens) and potential sires (DPQs, drone-producing queens) were fully phenotyped, tested on both the early and the late trait performance. Both dams and sires were selected and used for reproduction at 2 years old. In contrast to BQs, DPQs only genetically contribute drones to the next generations of breeding animals. Drones are born from unfertilized eggs, and thus, DPQs are often open-mated as this reduces costs (for example in Basso et al. 2024; Kistler et al. 2024). Accordingly, in our simulations, DPQs were always open-mated to base drones. The Initialization phase lasted from year 1 to year 10 included. In year 1, 24 unrelated and non-inbred founder BQs were created, mated to base drones, and produced worker groups to form colonies. In year 2, these colonies were phenotyped. Each BQ then produced 18 offspring BQs and 18 DPQs, which were open-mated to base drones and candidates for selection. Before colony phenotyping, 25% of all new BQs and 25% of all new DPQs were randomly eliminated to simulate mortality. In year 3 and each subsequent odd year, new BQ and DPQ colonies were phenotyped on the early and late trait, followed by a second random elimination of 25% of all colonies. As detailed previously, selection was for various alternative breeding goals where the weights for both traits differed. Among the 243 surviving colonies in each path, the best 24 BQ and the best 24 DPQ colonies were subjected to directional selection on a weighted sum of the phenotypes of both traits in standard deviation units. Each selected BQ produced 18 BQs and 18 DPQs in year 4 and each subsequent even year until year 10. Each of these offspring BQs, as all subsequent ones, were inseminated by drones produced by a single DPQ among the selected ones of that year, closing the breeding population. Each DPQ contributed to an equal number of matings. The choice of which DPQ to mate with each new BQ was random. In contrast, DPQs were always open-mated to base drones.

#### Base breeding scheme

Following this Initialization, in Base, BQs born in year 10 were selected in year 12 by directional selection again, but this time based on the weighted sum of the estimated breeding values (EBVs) of both traits. Everything else remained the same as in the Initialization phase. BQs and DPQs were thus fully phenotyped and had a 2-year generation interval each. The last queens were born in year 30, after 10 generations of selection following the Base breeding scheme.

#### Alternative breeding scheme

In parallel and starting from the same Initialization outcomes, in Alt, BQs born in year 10 were selected in year 11 at 1 year old. As a consequence, phenotyping was limited to the early trait only. As in Base, these dams were selected based on EBVs to produce offspring BQs and DPQs. In this transition year 11 from the Initialization to the Alt scheme, the offspring BQs were mated by the same pool of selected DPQs (born in year 8) as BQs born in year 10, due to the lack of DPQs born in year 9 in the initialization phase. After year 11, BQs were always mated by 2-year-old DPQs, which were fully phenotyped. Due to earlier selection, BQs in Alt faced only one annual mortality event before selection, resulting in a lower selection percentage of 7.4%, compared to 9.9% in Base, which was also the selection percentage for DPQs in both schemes. The last queens were born in year 30 as in Base, but corresponding here to 20 dam generations following the Alt breeding scheme.

#### Genetic connection across apiaries

Newborn BQs and DPQs were allocated to 18 apiaries in two independent draws for each queen type before the first mortality event. The resulting genetic connections between apiaries were as follows (we indicate average remaining numbers after mortality in brackets): each sister group was split on 3 apiaries, and each apiary tested 4 sister groups, each comprising 6 (4.5) queens. Each apiary thus tested 24 (18) BQs and 24 (18) DPQs. This resulted, on average, in 36 surviving colonies for phenotyping per apiary and test year.

#### Additional scenarios with increased selection percentages

In addition to the focal set of simulations, additional ones were run with an increased number of parents selected to produce the new generation (36 selected BQs and 36 selected DPQs). The total candidate population size and the number of apiaries were kept constant. Therefore, the selected proportions were increased on both dam and sire paths. Sister-group sizes were reduced from 18 to 12 potential queens on each path. Everything else remained the same. Results are only presented in the Tables S7 to S6.

### Genetic parameter sets and apiary effects

The residual and additive genetic variances in the base population were the same for both the early and the late trait.

The queen effect variance 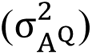 was half that of the worker effect 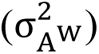 and a third of the residual effect 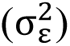.

These ratios of variances are in the range of real dataset estimates for most traits of interest, such as honey yield, royal jelly yield, swarming drive, and hygienic behavior (Brascamp *et al*., 2016, 2018; Hoppe *et al*., 2020; Basso *et al*., 2024).

Results will focus on simulations using a null genetic correlation between worker and queen effects (r_W,Q_). However, as often a negative r_W,Q_ is estimated in real datasets, a complete set of results for r_W,Q_ = −0.5 is also given in supplementary material.

Lastly, the genetic correlation between the early and the late (r_T1,T2_) trait was let to vary from [-0.6, +0.6] by 0.3 intervals. The resulting (co)variance matrix for both effects and traits was calculated as the following Kronecker product:

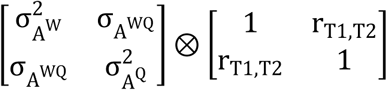

with 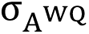 the genetic covariance between worker and queen effects.

Scenarios were defined by the combination of all breeding goals and all r_T1,T2_ values.

Simulated apiary effects for each year were drawn independently from a normal distribution. For the early and late trait, the variances of these apiary effects were set to represent approximately 1/3 and 2/3 of the phenotypic variance, respectively. These values are as estimated by Kistler *et al*. (2024), taking hygienic behavior and honey yield as examples of possible early and late traits.

### Estimation of genetic parameters and breeding values

Before each selection step in both Base and Alt, genetic parameters were estimated using a ReML approach applied to a 2-trait mixed linear model with worker and queen genetic effects (colony model) (Chevalet and Cornuet, 1982; Bienefeld *et al*., 2007) and a fixed environmental effect for the apiary by year effects.

All available phenotypes at the moment of selection were used to run the genetic evaluation, including in the performance file the phenotypes of colonies in the Initialization phase. This amounted to a minimum of 3264 records on each trait at the first estimation in Base (in year 12), and 2616 records on the late trait and 2940 on the early trait at the first estimation in Alt (in year 11). Note that in Alt, the early trait of 1-year-old DPQs was not included in the performance file, appearing only the following year together with the late trait, when these DPQs were 2 years old and became candidates to selection. In the last year, the performance file contained 9096 records for each trait in Base, while in Alt, it contained 8772 for the late trait and 15252 for the early trait.

The pedigree was completed assuming that each open-mated queen (BQs in years 1 and 2, and all DPQs) was mated to drones produced by a different pseudo sire composed of 100 unknown and unrelated dummy DPQs (see (Kistler *et al*., 2024). Later BQs, which were inseminated, had the single DPQ producing the drones mating them assigned as mate. This pedigree was used to obtain the inversed relationship matrix (**A**^−1^), containing a line and column for each dummy open mating pseudo sire, queen, and its worker group, following Brascamp and Bijma (2014, 2019b). Software to produce **A**^−1^ is available (Brascamp and Bijma, 2019).

The performance file and **A**^−1^were used as input for AIREMLF90 (ver. 1.149) of the BLUPF90 package (Misztal *et al*., 2002) to solve the BLUP equations. Starting parameter values for the convergence algorithm were set as, respectively, 0.9 and 1.1 times the true value for genetic and for residual variances. The convergence criterion was set to 1*e*^−11^. For each scenario, 50 to 55 independent replicate simulations were run. If convergence failed in either Base or Alt at any estimation, simulation stopped, recorded the fail, and a new replicate was run.

### Summary statistics

The genetic gain and inbreeding levels in the Alternative or Base breeding schemes were expressed compared to their levels for queens born in year 10 (end of Initialization).

To compare Alt and Base, relative differences were calculated as:

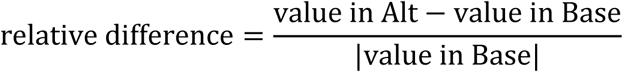

Results will show the relative differences calculated within repetitions and averaged across repetitions.

Alt’s empirical probability of bringing more genetic gain for the breeding goal than Base (p^) was estimated in each scenario as the percentage of replicated simulations in which Alt achieved strictly higher gains than Base. The standard error of this binomial variable was estimated as:

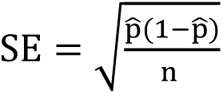, with n the num of replicated simulations.

## Results

Across all scenarios, convergence at each ReML estimation in both Base and Alt was reached for at least 80% of repetitions. Parameters most impeding convergence were a null r_W,Q_ and strong |r_T1,T2_| values.

The increase in breeding values (Fig. 2) and inbreeding was near linear after the Initialization phase, so that results at mid-term (after 10 years of selection following the Base or Alt breeding scheme) can be approximately inferred from results at long-term (20 years). Results will focus on the long-term outcomes.

**Fig. 2.**
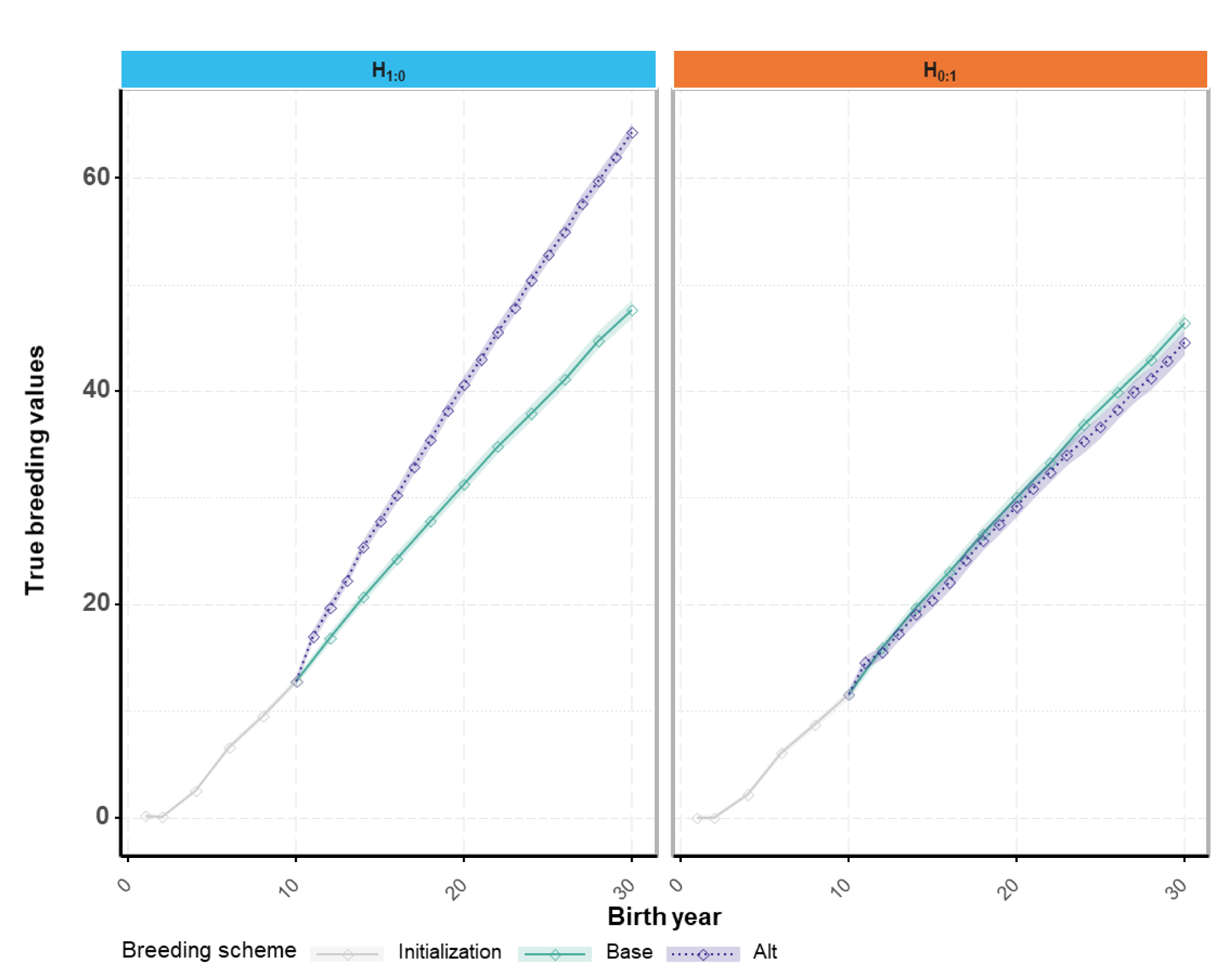
Genetic gain of potential dams across years of birth for selection on either the early or late trait and a null correlation between traits. H_1:0_: breeding goal comprising only the early trait H_0:1_: breeding goal comprising only the late trait Alt: accelerated breeding scheme in which only the early trait is phenotyped on potential dams, while potential sires are also phenotyped on the late trait. Partial phenotyping of the dams enables halving the dam generation interval to 1 year. Base: reference breeding scheme will full phenotyping and a 2-years generation interval on both the dam and the sire path.

### Genetic gain

#### Relative difference between Alt and Base in long-term genetic gains

Across all scenarios, Alt outperformed or equaled the average genetic gain obtained in Base except for the scenario focusing only on the late trait (H_0:1_) when r_T1,T2_ was null (Fig. 2, right pane). The averaged relative difference between Alt and Base in the genetic gain for the breeding goal went from -5% to +53% (values on top of the stacked bars in Fig. 3). Alt’s superiority in genetic gain was highest when the breeding goal focused only on the early trait (Fig. 2, left pane and Fig. 3), and it diminished as more weight was placed on the late trait. Alt brought similar relative gains compared to Base, regardless of r_T1,T2_, when the breeding goal focused on the early trait. This changed when more emphasis was placed on the late trait (H_0.5:0.5_ and H_0.25:0.75_), where Alt brought greater gains for positive r_T1,T2_ values than for null or negative values. In the extreme case where all the weight was on the late trait (H_0:1_), genetic gains in Alt relative to Base were equal for equal |r_T1,T2_| values, with larger |r_T1,T2_| being more beneficial to Alt.

**Fig. 3.**
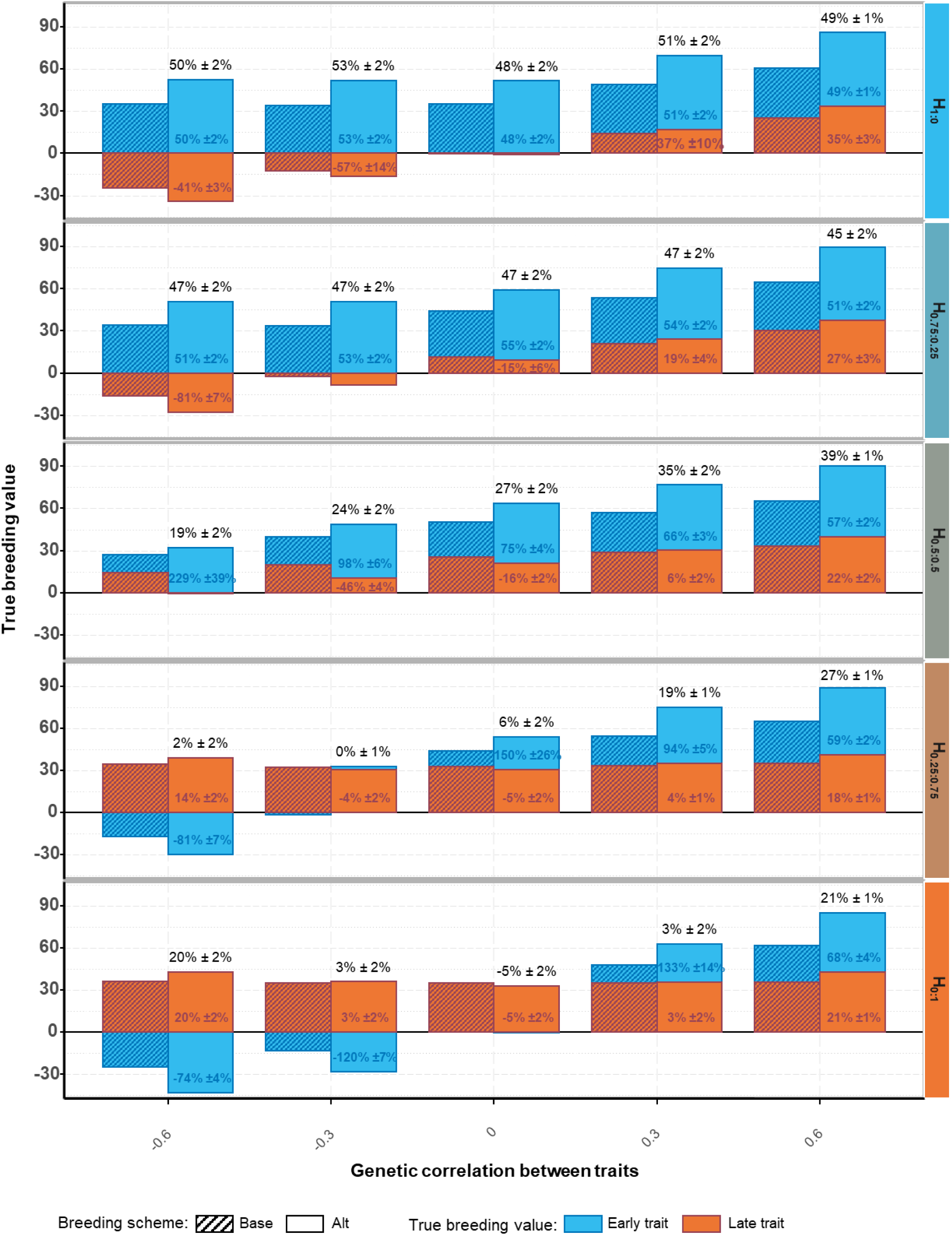
Increase in mean true breeding values and their relative difference between Alt and Base per trait and for the breeding goal after 20 years. H: breeding goal. The first number in the subscript refers to the weight on the early trait and the second number the weight on the late trait. Alt: accelerated breeding scheme in which only the early trait is phenotyped on potential dams, while potential sires are also phenotyped on the late trait. Partial phenotyping of the dams enables halving the dam generation interval to 1 year. Base: reference breeding scheme will full phenotyping and a 2-years generation interval on both the dam and the sire path. Values shown represent the relative differences between the outcomes of Alt and Base, calculated within repetitions and averaged across repetitions. These averaged relative differences were calculated for each trait (within bars) or for the breeding goal (above the stacked bars).

When worker and queen effects were negatively correlated (see Fig. S1) genetic gain decreased in both breeding schemes, but more so in Alt than in Base for breeding goals with equal or more weight on the late trait (H_0.5:0.5_, H0_.25:0.75_, and H_0:1_).

Fig. 3 also illustrates that the contribution of the two traits to the genetic gain for the breeding goal varies greatly between the two breeding schemes across scenarios. For example, for H_0.5:0.5_ and a null r_T1T2_, Alt outperformed by +27% Base for the gain on the breeding goal only due to its superior progress on the early trait (+75%), while it performed less well than Base for the late trait (-16%).

Lastly, the average relative difference between Alt and Base came from different gains in worker and queen effects across the breeding goals (Fig. S1 and S2). When the breeding goal emphasized the early trait, the superior genetic gain in Alt came from a mere balanced superiority in responses for the worker and queen effects in comparison to those observed with Base. However, as the breeding goal put more weight on the late trait, Alt’s superiority was achieved through higher gains for queen effects than for worker effects relative to Base. This was due to Alt achieving less gains on worker effects as the breeding goal put more weight on the late trait, rather than to Alt increasing its gains on queen effects. For example, for H_0.25:0.75_ and r_T1,T2_ = 0, Alt brought 7% less genetic gain on worker effects, but this was more than compensated by a +36% relative increase on queen effects, resulting in a +6% gain on the breeding goal compared to Base.

#### Probability of the Alternative breeding scheme to outperform the Base breeding scheme

Table 1 shows the empirical probability of Alt to outperform Base in terms of genetic gain for the breeding goal and for each trait.

**Table 1:** Empirical probability of the Alternative breeding scheme to bring more genetic gain than the Base one after 20 years of selection.

| Breeding goal (H) | $r_{T1,T2}$ | Empirical probability of the Alt breeding scheme to bring more genetic gain than the Base one | | |
| --- | --- | --- | --- | --- |
|  |  | for H | for T1 | for T2 |
| **H<sub>1:0</sub>** | 0 | 1.00 (NE) | 1.00 (NE) | 0.48 (0.07) |
|  | -0.6 | 1.00 (NE) | 1.00 (NE) | 0.02 (0.02) |
|  | 0.6 | 1.00 (NE) | 1.00 (NE) | 0.98 (0.02) |
| **H<sub>0.75:0.25</sub>** | 0 | 1.00 (NE) | 1.00 (NE) | 0.32 (0.06) |
|  | -0.6 | 1.00 (NE) | 1.00 (NE) | 0.00 (NE) |
|  | 0.6 | 1.00 (NE) | 1.00 (NE) | 0.98 (0.02) |
| **H<sub>0.5:0.5</sub>** | 0 | 0.98 (0.02) | 1.00 (NE) | 0.14 (0.05) |
|  | -0.6 | 0.90 (0.04) | 1.00 (NE) | 0.00 (NE) |
|  | 0.6 | 1.00 (NE) | 1.00 (NE) | 0.98 (0.02) |
| **H<sub>0.25:0.75</sub>** | 0 | 0.68 (0.07) | 0.94 (0.03) | 0.28 (0.06) |
|  | -0.6 | 0.62 (0.07) | 0.02 (0.02) | 0.86 (0.05) |
|  | 0.6 | 1.00 (NE) | 1.00 (NE) | 1.00 (NE) |
| **H<sub>0:1</sub>** | 0 | 0.26 (0.06) | 0.50 (0.07) | 0.26 (0.06) |
|  | -0.6 | 0.94 (0.03) | 0.00 (NE) | 0.94 (0.03) |
|  | 0.6 | 0.98 (0.02) | 1.00 (NE) | 0.98 (0.02) |

NE stands for non-estimable.

When all or most the weight in the breeding goal was on the early trait, the empirical probability of Alt to bring more genetic gain for the breeding goal than Base was 1, and at least 0.9 when weights on each trait in the breeding goal were equal. Shifting the emphasis to the late trait, this probability decreased to 0.62 for H_0.25:0.75_ and a null r_T1,T2_. Still for a null r_T1,T2_ but when all the weight was on the late trait, the probability for Alt to bring more genetic gain on the breeding goal than Base was only 0.26, this scenario being the worst for Alt.

### Inbreeding

#### Long-term inbreeding level

The increases in mean inbreeding level reached on the long term were similar across all scenarios for both breeding schemes (Fig. 4 and Table S1). They averaged 23% in Base and 28% in Alt for r_W,Q_ = 0. In Base, it reached its maximum at 24% for H_0.5:0.5_ with the strongest negative r_T1,T2_. The increases were higher in Alt by 17 ± 3% to 31 ± 3% compared to Base (+24% on average), with larger values for breeding goals emphasizing the late trait. The increase in mean inbreeding level in Alt reached a maximum value of 30% for H_0:1_ and r_T1,T2_ = 0 (Table 2).

**Fig. 4.**
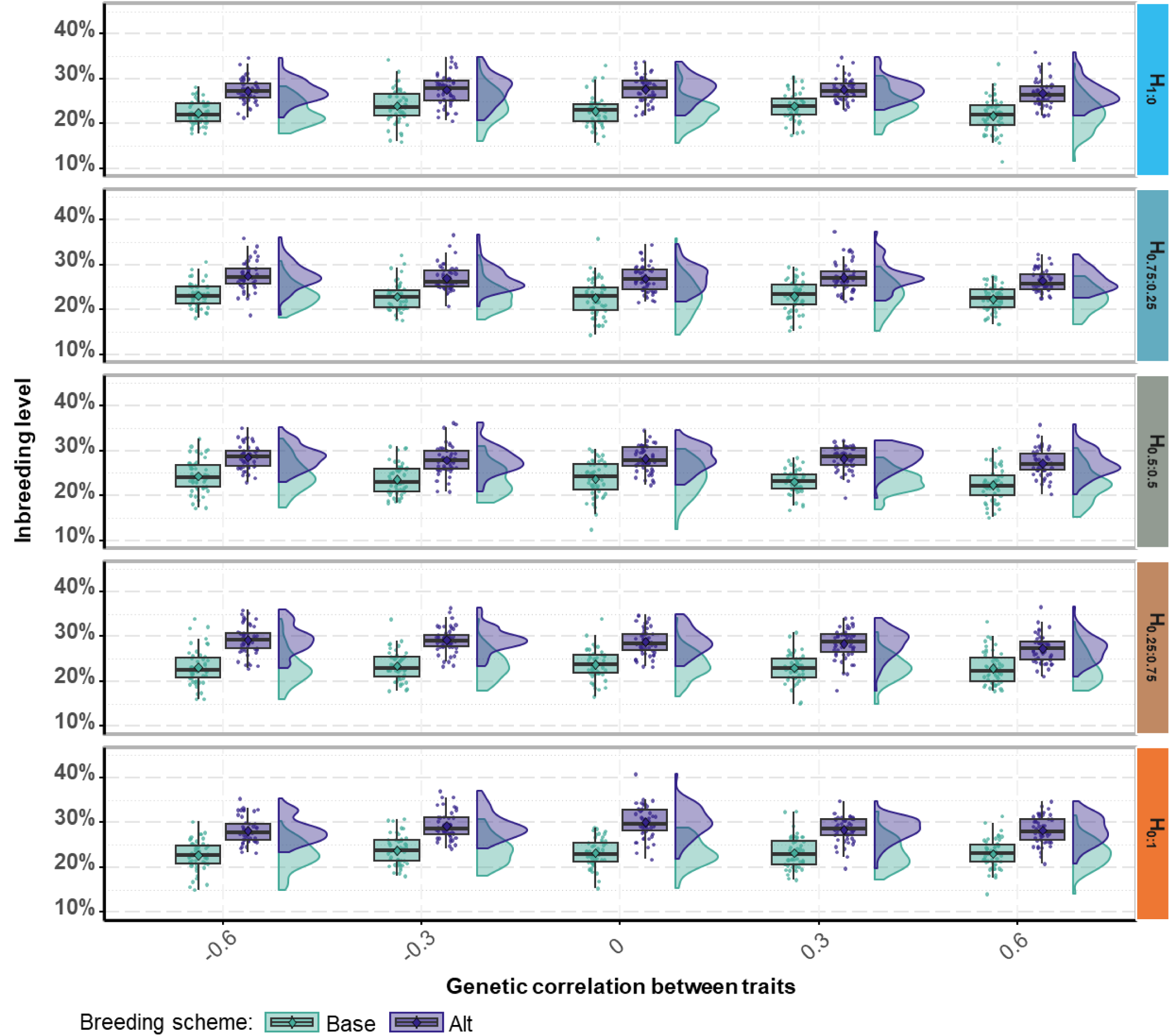
Increase in inbreeding level in the Base and Alt breeding scheme for all scenarios with a null between-effects genetic correlation. H: breeding goal. The first number in the subscript refers to the weight on the early trait and the second number the weight on the late trait. Alt: accelerated breeding scheme in which only the early trait is phenotyped on potential dams, while potential sires are also phenotyped on the late trait. Partial phenotyping of the dams enables halving the dam generation interval to 1 year. Base: reference breeding scheme will full phenotyping and a 2-years generation interval on both the dam and the sire path.

**Table 2:**
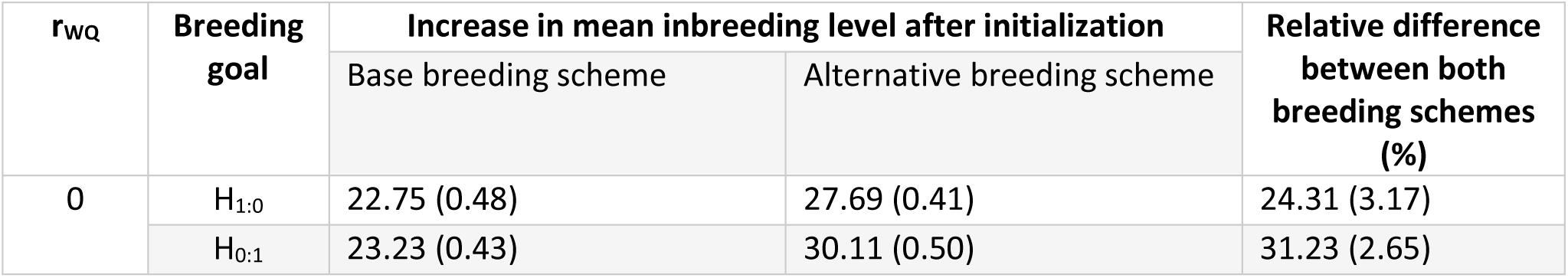
Increase in mean inbreeding level (%) after initialization and its relative difference between Base and Alt for the two single-trait breeding goals with a null genetic correlation between traits

Results for the same scenarios but with r_W,Q_ = −0.5 were very similar, with the same or a maximum of +2% of added inbreeding levels (Table S2).

## Discussion

We used simulation to quantify the impact of reducing the classical 2-years dam generation interval in honeybee breeding schemes to 1 year. This shortening in the dam’s generation interval implied a partial phenotyping limited to some early recorded traits. The sire generation interval remained at 2 years with phenotyping of both early and late traits. The effect of various breeding goals was studied, emphasizing either an early or a late trait or equally weighting both. Different genetic correlations between both traits were also considered.

Annual genetic gain for the breeding goal was generally greater with the shorter generation interval strategy than with the base scenario with its longer one. This greater gain was achieved more through the gain in the early trait compared to the gain in the late trait. It came, however, at the cost of an increase in mean inbreeding level of roughly +20% to +30% after 20 years of selection.

### Major impact of the dam index accuracy on the differences in relative gain between Alt and Base across breeding goals

Regardless of the breeding goal, response to selection was favored in Alt due to a 25% reduction in the average generation interval and a 7.5% increase in dam selection intensity, as earlier dam selection avoided a second random mortality event. Still regardless of the breeding goal, the sire index accuracies were roughly similar in both breeding schemes as phenotypes for both traits were available directly on candidates in Alt as in Base (see Fig. S3, left panel), small differences being probably due to differences in available information on relatives and in inbreeding levels. On the opposite, dam index accuracies varied widely across breeding goals in Alt, explaining most of the differences in genetic gain compared to Base.

The dam index accuracies ranged from +6% to -63% (see Fig. S3, right panel) in Alt compared to Base, depending on the breeding goal. The stronger reduction in accuracy arose for breeding goals emphasizing the late trait, as the late trait was not phenotyped on candidate dams in Alt. Conversely, when the breeding goal focused only on the early trait, the dam index accuracies were similar in Base and Alt, as candidate dams were always phenotyped on the early trait. We analyzed dam index accuracies in Alt using selection-index theory (Hazel, 1943) for a two-trait breeding goal with only one trait used as selection criterion. Simulations confirmed the expected theoretical patterns: when both traits had equal weights, index accuracy increased with r_T1,T2_. When all the weight is on the unmeasured trait, the accuracy depends on 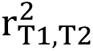, making it independent of the correlation’s sign. In the intermediary case of unbalanced weights, the index accuracy depends of both r_T1,T2_ and 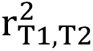, reaching the lowest values at moderate negative correlations and the highest at strong positive correlations.

While maintaining early dam selection, candidate dams could be phenotyped after selection to increase the accuracy in Alt for breeding goals emphasizing the late trait. By recording the late trait for older queens, these data would be integrated in the EBVs of the parents and, consequently, in the EBVs of the new candidate dams at the moment of selection. Doing so in simulation for H_0:1_ and null values for r_T1,T2_and r_WQ_ increased the dam selection accuracy from 0.21 to 0.28, resulting in a 21% increase in genetic gain. Even in this least favorable scenario to Alt, more genetic gain could then be achieved using this parental information than with Base (+17% compared to -5% when these late-trait records on older candidate dams were absent, data not shown). If the phenotyping of late traits comes at a reduced cost, as arguably for total honey yield, later recording might thus be of interest to get a more efficient breeding scheme.

### Inbreeding

The Alt breeding scheme appeared to be a favorable strategy for genetic gain but unfavorable for inbreeding per year, reaching +17 ± 3% to +31% ± 3% higher mean inbreeding levels. The highest difference in inbreeding between the two breeding schemes was observed for a null r_T1,T2_and when the breeding goal focused on the late trait, which was not phenotyped on candidate BQs. In the absence of phenotypes on candidates, using BLUP-EBVs, all sister- queens had similar EBVs for the target trait, corresponding to the mean of their parents’ EBVs (they still differed due to the fact that the between trait correlation was estimated and not fixed to 0). This probably resulted in the selection of BQs originating mostly from only the few best sister groups, increasing inbreeding more rapidly than when better discrimination within sister-groups could be made. Across scenarios, the general higher inbreeding levels in Alt was mainly due to Alt’s 33% increase in the average number of generations compared to Base during the same period, the inbreeding rate being actually reduced in Alt by -2% ± 2% to -12% ± 2% per generation (-8% on average, Table S3 and S4).

However, in both breeding schemes and for all breeding goals and between-traits correlations, inbreeding rates were above two times the common recommended maximum 1% rate (FAO, 2013) per generation. It might be noticed, however, that breeding schemes in reality, especially in honeybees, often do not remain genetically strictly closed for decades. However, maintaining the population closed and its size fixed, inbreeding rates could be lowered by relaxing the selection intensity, such as by selecting more breeding queens as in our additional simulations with 36 selected BQs and DPQs per generation (instead of 24, representing 33% less selected queens). This strategy effectively reduced inbreeding rates (near but slightly under the expected -33% reduction, S5 and S6), while reducing genetic gains by only -5% on average across all scenarios whatever the r_WQ_ value (Table S7 and S8). In particular, genetic gain was at most only marginally reduced in Alt with 36 BQs, while the increased number of selected queens reduced mean inbreeding rates by 24% ± 2% to 33% ± 2% (Table S5 and S6). Alt with 36 BQs thus led to at least almost as much genetic gain than Base with 24 BQs (-2%

± 2% for H_0:1_ and r_T1,T2_ = 0) or significantly better (around +41% ± 2% for H_1:0_ whatever the r_T1,T2_), while also bringing less long-term inbreeding levels (-5% for H_0:1_ and r_T1,T2_ = 0, to - 18% for H_1:0_ & r_T1,T2_ = −0.6). This shows that if a scheme similar to Alt is used in practice, maintaining a sufficient population size becomes more important to limit inbreeding development in time due to the accelerated generation turnover. The relaxed intensity can come at practically no costs in genetic gain benefitting from the increased genetic variance remaining, while the reduced generation interval enables benefitting from the greater genetic gains per year.

Another approach to relax selection intensity in order to maintain genetic diversity could have been to perform within-family selection (Kistler *et al*., 2021). However, an optimal way to compare Alt and Base for mean genetic gains at equal levels of inbreeding would have been to use Optimal Contribution Selection (Meuwissen, 1997) to fix a predetermined inbreeding rate. Unfortunately, this method still has to be transposed to honeybees and their specificities, in particular the early mating before phenotyping, and therefore before selection, and haplo- diploïdy and polyandry impacting the different genetic contribution of queens when used as sires or dams.

### Reduced generation intervals and longevity

The concerning high and increasing colony losses in honeybees are partly due to issues in queen quality and longevity (Amiri *et al*., 2017). Our study simulated colony mortality as a fully random event between two years. In reality, however, survival might not be entirely random, and longer generation intervals might reveal genetic differences in longevity. By shortening generation intervals, there would be no chance of indirect selection on longevity.

### Splitting the population in two when having a two-year generation interval

To have the most straight-forward comparison, the only difference between Alt and Base was the reduced dam generation interval. In consequence, in Alt, potential dams and sires were bred every year, as opposed to every even year in Base. In reality, breeders using a scheme similar to Base would have probably split their population in two, to breed new candidates in one subpopulation and phenotype the other every year, enabling them to increase the total population size and to cross-breed excelling individuals from one population with the other (Uzunov *et al*., 2022). This would likely benefit genetic gain and inbreeding.

### Shortening of the generation interval at no expense of phenotyping late traits

Breeders could potentially benefit from the accelerated response to selection obtained by the shortening of the dam generation interval without having to sacrifice recording of late traits before selection. This could be achieved by delaying selection, reproduction, and mating of new candidate dams to after the period of measurement of late traits. In temperate climates this might be achieved by mating new potential dams after the production season, in late summer or autumn. Depending on the region, colonies might accept more or less easily to produce drones in that period. Alternatively, semen might be collected earlier when drones can reliably be obtained from selected sires and stored to be used in insemination a few months later (van Praagh *et al*., 2014; Hopkins *et al*., 2017).

### Conclusion

Halving the dam generation interval to one year at the expense of phenotyping only the early trait on potential dams generally resulted in an increased genetic gain, up to +50% after 20 years of selection depending on the breeding goal. The highest increases were observed when the breeding goal focused primarily on the early trait or when the genetic correlation between the early and late traits was positive. In contrast, the accelerated breeding scheme achieved slightly lower genetic gain than the Base one when the breeding goal focused only on the late trait and when it was not correlated with the early recorded trait. Although inbreeding rates per generation were slightly reduced in the accelerated breeding scheme, inbreeding levels were still higher due to the increase in the number of generations obtained during the 20 years of selection. Shortening the generation interval should therefore be accompanied by strategies to limit long-term inbreeding. When accelerating the breeding scheme, an effective strategy is to increase the breeding nucleus size by relaxing selection intensity. Doing so can substantially reduce inbreeding rates while still retaining most of the genetic gain benefits compared to not shortening the average generation interval.

## Supporting information

Supplementary file 1 and all supplementary tables

## Acknowledgements

The main author is thankful to all the partners involved in the research projects linked to this work, in particular Coline Kouchner and the beekeepers involved in the beekeeping association ADAPI, as well as beekeepers from Melyos Apicoltura.

The authors wish also to extend their acknowledgments to the funders that supported this work via a public grant overseen by the French National Research Agency (ANR) as part of the « Investissements d’Avenir » program, through the "ADI 2021" project funded by IDEX Paris-Saclay (ANR-11-IDEX-0003-02) and also by CARNOT France Future Elevage (project BeeMuSe).

## Author contribution statement

TK developed the R code for the simulations, performed the simulations and analyzed the data generated, and was involved in the conception of the work, the interpretation of results, and wrote the original draft. BB was involved in the conception of the work. PB, EWB, and FP supervised the work and were involved in its conception, methodological choices, interpretation of the results and revisions to the draft. All authors read and approved the final manuscript.

## Data archiving

The R simulation code used in the current study has been made publicly available at the simu_bees GitHub repository https://github.com/Tristan-Kistler/simu_bees in the folder ‘partial_pheno_short_gen_interval_study’ and archived on Software Heritage, with the SWHID: swh:1:dir:4f183d1233df180713db7ef3395cbbb07a4e01f0;origin=Kistler/simu_bees;visit=swh:1:snp:7b9d55a2a2a84e37152a47143857d58a37eb2c82;anchor=swh:1:rev:afed68dfd1d6a0691a21f99479dd2fc9e28de036;path=/partial_pheno_short_gen_inte rval_study/https://github.com/Tristan-Kistler/simu_bees;visit=swh:1:snp:7b9d55a2a2a84e37152a47143857d58a37eb2c82;anchor=swh:1:rev:afed68dfd1d6a0691a21f99479dd2fc9e28de036;path=/partial_pheno_short_gen_interval_study/.

Further developments will also be available via the GitHub repository.

The R version used to run the above code was 3.5.0 (2018-04-23), but should be compatible with versions from 3.6.2 to 4.2.1.

## Research Ethics Statement

Not applicable.

---

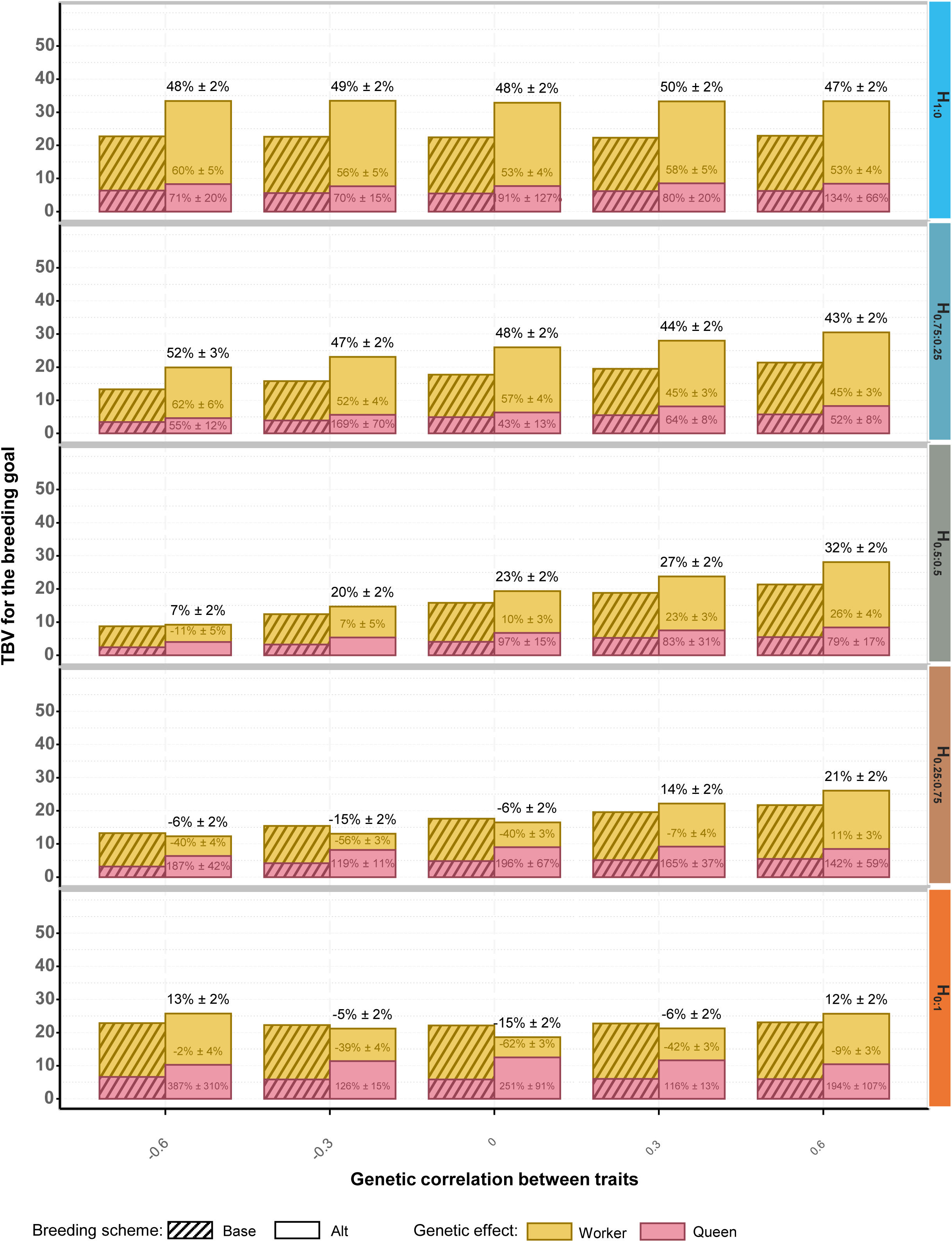

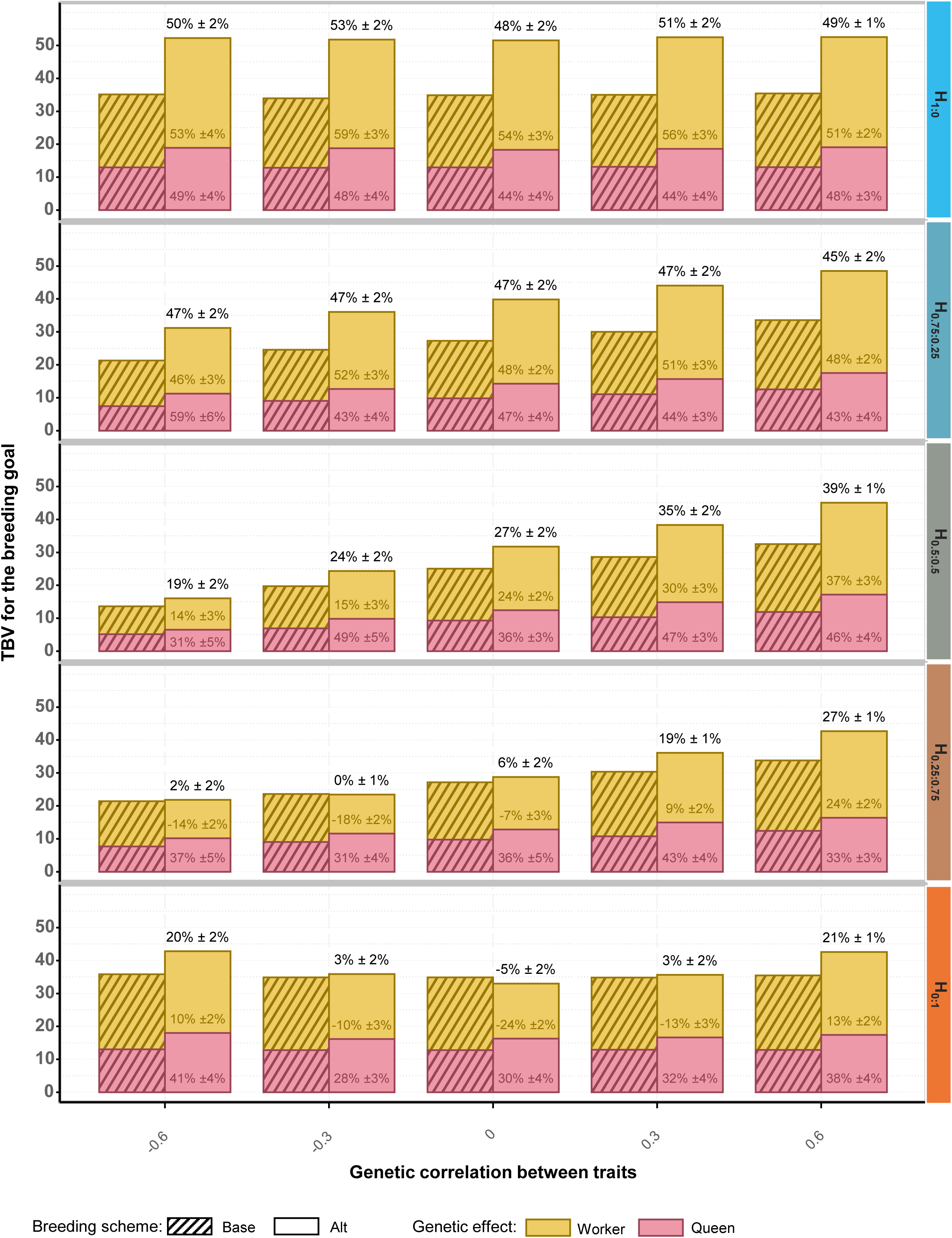

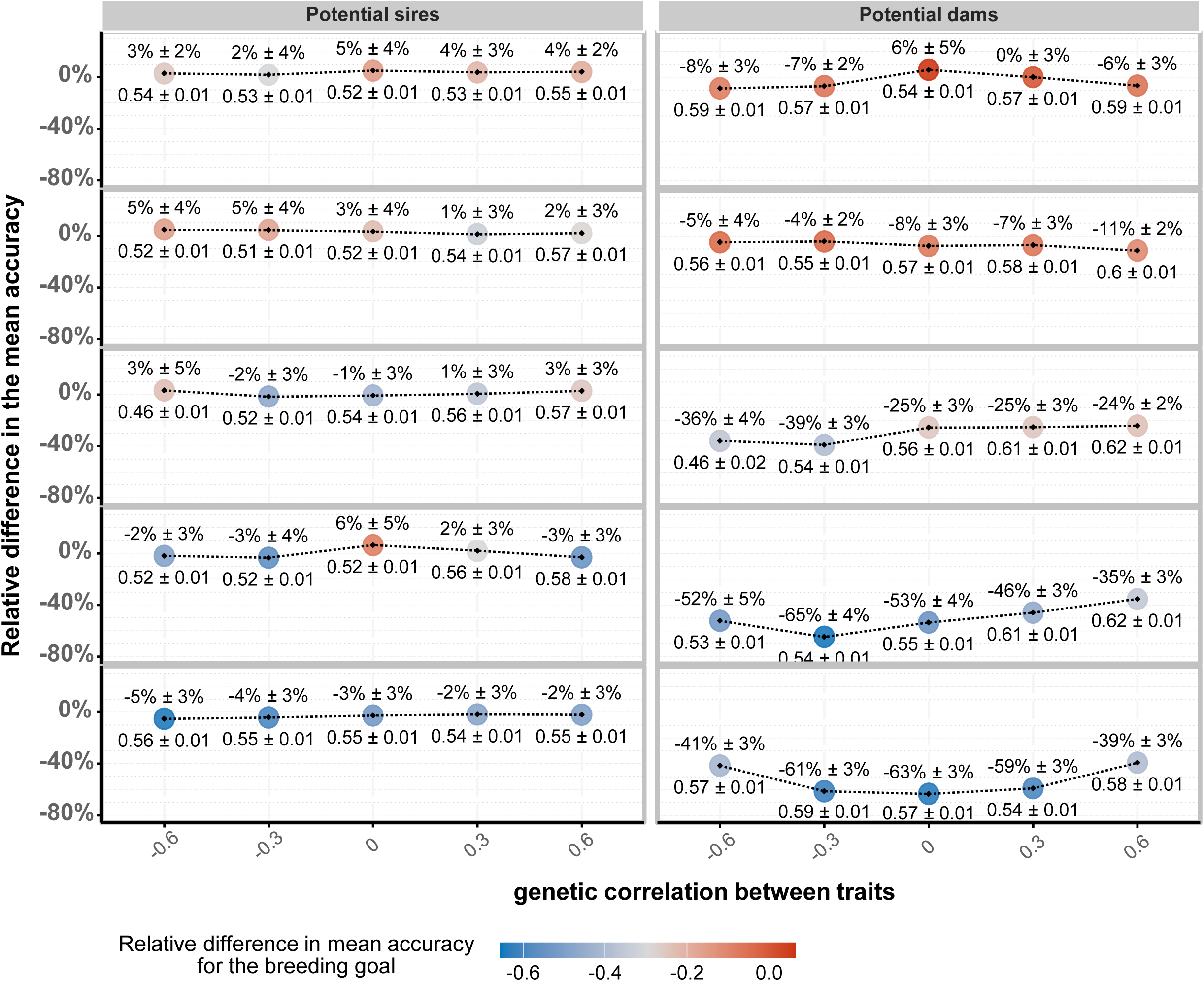

