## Supplementary file 1 and all supplementary tables for "How partial phenotyping to reduce generation intervals can help to increase annual genetic gain in selected honeybee populations"

**File S1**

**Infinitesimal model applied to the honeybee for queens, drones, and worker groups**

Generation of base individuals:

$\mathbf{a}^{\mathbf{base\ BQs}} \sim \boldsymbol{N}(\mathbf{0}, \boldsymbol{\Sigma}^2_{\mathbf{a}})$: Breeding queens (BQ) from the base population had their additive genetic values **(a)** drawn from a bivariate normal distribution centered on **0** and with (co)variance matrix of both effects and traits $\boldsymbol{\Sigma}^2_{\mathbf{a}}$.

$\mathbf{a}^{\mathbf{base\ Ds}} \sim \boldsymbol{N}(\mathbf{0}, \tfrac{1}{2}\boldsymbol{\Sigma}^2_{\mathbf{a}})$: Base drones, which are haploid, had their additive genetic values made to (co)vary half as much as that of BQs.

After mating, queens produced offspring worker groups by:

$$\overline{\mathbf{a}^{\mathbf{W}}} = \frac{1}{2} \cdot \mathbf{a}^{\mathbf{Q}} + \overline{\mathbf{a}^{\mathbf{Ds}}}$$

where the additive genetic values of a worker group ($\overline{\mathbf{a}^{\mathbf{W}}}$) equaled the sum of half the additive genetic values of its BQ and the average additive genetic values of the drones that mated the BQ.

Colonies' phenotypes were obtained using:

$$\mathrm{y} = \mathrm{a}_{\mathrm{q}}^{\mathrm{Q}} + \frac{1}{2}\mathrm{a}_{\mathrm{w}}^{\mathrm{Q}} + \overline{\mathrm{a}_{\mathrm{w}}^{\mathrm{Ds}}} + \mathrm{E} + \varepsilon$$

where the performance (y) of a colony equaled the sum of the queen genetic effect expressed by the queen $\left(\mathrm{a}_{\mathrm{q}}^{\mathrm{Q}}\right)$, half the worker genetic effect of the queen $\left(\frac{1}{2}\mathrm{a}_{\mathrm{w}}^{\mathrm{Q}}\right)$, the average worker genetic effect of the drones that mated the queen, a non-heritable apiary by year effect (E), and a random residual (ε) modeling other non-additive genetic effects .

Offspring queens inherited their additive genetic values following:

$$\mathbf{a}^{\mathbf{Q}} = \frac{1}{2} \cdot \mathbf{a}^{\mathbf{BQ}} + \mathbf{a}^{\mathbf{D}} + \boldsymbol{\varphi}^{\mathbf{BQ}}$$

where their additive genetic values were the sum of half that of their BQ's, that of one of the drones that mated their BQ, and a mendelian sampling term ($\boldsymbol{\varphi}$) drawn from $\boldsymbol{N}(\mathbf{0}, \frac{1}{4}\cdot(1\text{-F})\cdot\boldsymbol{\Sigma}^2\mathbf{a})$, where F is the inbreeding coefficient of the offspring's BQ.

Lastly, drones inherited their additive genetic values following:

$$\mathbf{a}^{\mathbf{D}} = \frac{1}{2}\mathbf{a}^{\mathbf{DPQ}} + \boldsymbol{\varphi}^{\mathbf{DPQ}} \quad (6)$$

where haploid drones' additive genetic values were inherited only from their drone-producing queen (DPQ).

Further details are given in Kistler et al. [20].

*Inbreeding level*

**Table S1: Increase in mean inbreeding level (%) after initialization and relative difference in the Base and Alt breeding scheme for all scenarios with a null between-effects genetic correlation.**

| Breeding goal | $r_{T1,T2}$ | Added inbreeding level (%) | | Relative difference between both breeding schemes (%) |
|---|---|---|---|---|
| | | Base breeding scheme | Alt breeding scheme | |
| $H_{1:0}$ | 0 | 22.75 (0.48) | 27.69 (0.41) | 24.31 (3.17) |
| | -0.3 | 23.97 (0.54) | 27.49 (0.47) | 17.23 (3.09) |
| | -0.6 | 22.42 (0.37) | 27.24 (0.37) | 22.77 (2.28) |
| | 0.3 | 23.96 (0.44) | 27.60 (0.35) | 17.20 (2.74) |
| | 0.6 | 21.79 (0.55) | 26.74 (0.43) | 25.93 (3.31) |
| $H_{0.75:0.25}$ | 0 | 22.56 (0.55) | 26.83 (0.40) | 22.12 (3.01) |
| | -0.3 | 22.91 (0.43) | 26.94 (0.40) | 19.13 (2.44) |
| | -0.6 | 23.12 (0.38) | 27.49 (0.44) | 20.09 (2.42) |
| | 0.3 | 22.95 (0.48) | 27.20 (0.46) | 20.75 (2.84) |
| | 0.6 | 22.39 (0.39) | 26.47 (0.35) | 19.97 (2.54) |
| $H_{0.5:0.5}$ | 0 | 23.72 (0.55) | 28.20 (0.40) | 21.78 (3.03) |
| | -0.3 | 23.62 (0.48) | 27.92 (0.45) | 20.34 (2.80) |
| | -0.6 | 24.32 (0.51) | 28.51 (0.38) | 19.57 (2.79) |
| | 0.3 | 23.05 (0.36) | 28.27 (0.36) | 23.91 (2.27) |
| | 0.6 | 22.32 (0.52) | 27.21 (0.43) | 24.46 (2.92) |
| $H_{0.25:0.75}$ | 0 | 23.72 (0.46) | 28.80 (0.41) | 23.63 (2.98) |
| | -0.3 | 23.49 (0.46) | 29.21 (0.39) | 26.50 (2.80) |
| | -0.6 | 23.17 (0.56) | 29.28 (0.46) | 29.16 (3.27) |
| | 0.3 | 23.02 (0.44) | 28.45 (0.47) | 25.12 (2.67) |
| | 0.6 | 22.98 (0.51) | 27.30 (0.43) | 21.69 (3.33) |
| $H_{0:1}$ | 0 | 23.23 (0.43) | 30.11 (0.50) | 31.23 (2.65) |
| | -0.3 | 23.80 (0.45) | 29.24 (0.42) | 24.53 (2.44) |
| | -0.6 | 22.67 (0.46) | 28.19 (0.40) | 26.96 (3.34) |
| | 0.3 | 23.25 (0.49) | 28.42 (0.39) | 24.64 (2.91) |
| | 0.6 | 23.03 (0.46) | 28.30 (0.44) | 24.58 (2.46) |

**Table S2: Increase in mean inbreeding level (%) after initialization and relative difference in the Base and Alt breeding scheme for all scenarios with a between-effects genetic correlation of -0.5.**

| Breeding goal | $r_{T1,T2}$ | Added inbreeding level (%) | | Relative difference between both breeding schemes (%) |
|---|---|---|---|---|
| | | Base breeding scheme | Alt breeding scheme | |
| $H_{1:0}$ | 0 | 24.38 (0.46) | 29.21 (0.45) | 21.74 (2.68) |
| | -0.3 | 24.56 (0.52) | 28.96 (0.41) | 20.29 (2.63) |
| | -0.6 | 23.64 (0.43) | 28.28 (0.38) | 21.38 (2.44) |
| | 0.3 | 23.73 (0.44) | 29.48 (0.46) | 26.11 (2.73) |
| | 0.6 | 23.54 (0.41) | 28.05 (0.37) | 20.99 (2.63) |
| $H_{0.75:0.25}$ | 0 | 23.78 (0.47) | 28.24 (0.39) | 20.91 (2.59) |
| | -0.3 | 23.92 (0.45) | 28.75 (0.43) | 22.19 (2.72) |
| | -0.6 | 23.35 (0.49) | 29.15 (0.38) | 27.27 (2.73) |
| | 0.3 | 24.41 (0.48) | 28.82 (0.34) | 20.31 (2.55) |
| | 0.6 | 23.33 (0.47) | 28.54 (0.44) | 24.04 (2.53) |
| $H_{0.5:0.5}$ | 0 | 24.31 (0.53) | 28.57 (0.40) | 20.28 (2.78) |
| | -0.3 | 24.30 (0.48) | 28.76 (0.42) | 20.64 (2.82) |
| | -0.6 | 25.15 (0.47) | 29.38 (0.39) | 18.66 (2.53) |
| | 0.3 | 24.35 (0.42) | 29.30 (0.44) | 21.59 (2.15) |
| | 0.6 | 23.82 (0.42) | 28.65 (0.50) | 21.55 (2.41) |
| $H_{0.25:0.75}$ | 0 | 23.93 (0.43) | 28.76 (0.33) | 21.80 (2.10) |
| | -0.3 | 23.71 (0.51) | 29.71 (0.34) | 28.43 (3.17) |
| | -0.6 | 24.44 (0.48) | 29.62 (0.44) | 23.16 (2.71) |
| | 0.3 | 24.02 (0.46) | 29.06 (0.52) | 22.72 (2.79) |
| | 0.6 | 24.24 (0.36) | 28.60 (0.43) | 19.13 (2.33) |
| $H_{0:1}$ | 0 | 23.45 (0.47) | 30.31 (0.38) | 32.00 (3.08) |
| | -0.3 | 24.15 (0.39) | 29.54 (0.39) | 24.07 (2.59) |
| | -0.6 | 24.26 (0.50) | 28.97 (0.38) | 21.74 (2.70) |
| | 0.3 | 24.20 (0.50) | 29.18 (0.37) | 22.96 (2.65) |
| | 0.6 | 24.07 (0.53) | 28.27 (0.43) | 19.75 (2.75) |

### *Inbreeding rate*

**Table S3: Inbreeding rate and relative difference in the Base and Alt breeding scheme for all scenarios with a null between-effects genetic correlation.**

| Breeding goal | $r_{T1,T2}$ | Inbreeding rate (%) | | Relative difference between both breeding schemes (%) |
|---|---|---|---|---|
| | | Base breeding scheme | Alt breeding scheme | |
| $H_{1:0}$ | 0 | 2.28 (0.05) | 2.08 (0.03) | -6.79 (2.38) |
| | -0.3 | 2.40 (0.05) | 2.06 (0.03) | -12.09 (2.32) |
| | -0.6 | 2.24 (0.04) | 2.04 (0.03) | -7.94 (1.72) |
| | 0.3 | 2.39 (0.04) | 2.07 (0.03) | -12.10 (2.05) |
| | 0.6 | 2.18 (0.05) | 2.01 (0.03) | -5.58 (2.48) |
| $H_{0.75:0.25}$ | 0 | 2.29 (0.06) | 2.02 (0.03) | -9.44 (2.32) |
| | -0.3 | 2.29 (0.04) | 2.02 (0.03) | -10.69 (1.83) |
| | -0.6 | 2.31 (0.04) | 2.06 (0.03) | -9.92 (1.81) |
| | 0.3 | 2.29 (0.05) | 2.04 (0.03) | -9.41 (2.12) |
| | 0.6 | 2.24 (0.04) | 1.99 (0.03) | -10.01 (1.90) |
| $H_{0.5:0.5}$ | 0 | 2.39 (0.05) | 2.11 (0.03) | -9.22 (2.27) |
| | -0.3 | 2.36 (0.05) | 2.09 (0.03) | -9.70 (2.10) |
| | -0.6 | 2.43 (0.05) | 2.14 (0.03) | -10.31 (2.10) |
| | 0.3 | 2.31 (0.04) | 2.12 (0.03) | -7.12 (1.72) |
| | 0.6 | 2.23 (0.05) | 2.04 (0.03) | -6.61 (2.19) |
| $H_{0.25:0.75}$ | 0 | 2.37 (0.05) | 2.16 (0.03) | -7.30 (2.24) |
| | -0.3 | 2.35 (0.05) | 2.19 (0.03) | -5.06 (2.11) |
| | -0.6 | 2.32 (0.06) | 2.20 (0.03) | -3.09 (2.45) |
| | 0.3 | 2.30 (0.04) | 2.14 (0.04) | -6.20 (1.99) |
| | 0.6 | 2.30 (0.05) | 2.05 (0.03) | -8.80 (2.50) |
| $H_{0:1}$ | 0 | 2.33 (0.04) | 2.26 (0.04) | -1.68 (1.99) |
| | -0.3 | 2.38 (0.05) | 2.19 (0.03) | -6.59 (1.83) |
| | -0.6 | 2.29 (0.05) | 2.11 (0.03) | -5.75 (2.42) |
| | 0.3 | 2.33 (0.05) | 2.13 (0.03) | -6.53 (2.18) |
| | 0.6 | 2.30 (0.05) | 2.12 (0.03) | -6.58 (1.84) |

**Table S4: Inbreeding rate and relative difference in the Base and Alt breeding scheme for all scenarios with a between-effects genetic correlation of -0.5.**

| Breeding goal | $r_{T1,T2}$ | Inbreeding rate (%) | | Relative difference between both breeding schemes (%) |
|---|---|---|---|---|
| | | Base breeding scheme | Alt breeding scheme | |
| $H_{1:0}$ | 0 | 2.44 (0.05) | 2.19 (0.03) | -8.68 (2.00) |
| | -0.3 | 2.46 (0.05) | 2.17 (0.03) | -9.77 (1.96) |
| | -0.6 | 2.36 (0.04) | 2.12 (0.03) | -8.92 (1.83) |
| | 0.3 | 2.37 (0.04) | 2.21 (0.03) | -5.43 (2.05) |
| | 0.6 | 2.35 (0.04) | 2.10 (0.03) | -9.22 (1.96) |
| $H_{0.75:0.25}$ | 0 | 2.37 (0.05) | 2.12 (0.03) | -9.30 (1.94) |
| | -0.3 | 2.39 (0.04) | 2.16 (0.03) | -8.36 (2.05) |
| | -0.6 | 2.33 (0.05) | 2.19 (0.03) | -4.50 (2.06) |
| | 0.3 | 2.44 (0.05) | 2.16 (0.03) | -9.76 (1.91) |
| | 0.6 | 2.33 (0.05) | 2.14 (0.03) | -6.89 (1.91) |
| $H_{0.5:0.5}$ | 0 | 2.43 (0.05) | 2.14 (0.03) | -9.82 (2.09) |
| | -0.3 | 2.43 (0.05) | 2.16 (0.03) | -9.50 (2.12) |
| | -0.6 | 2.52 (0.05) | 2.20 (0.03) | -11.01 (1.89) |
| | 0.3 | 2.43 (0.04) | 2.20 (0.03) | -8.81 (1.62) |
| | 0.6 | 2.38 (0.04) | 2.15 (0.04) | -8.84 (1.81) |
| $H_{0.25:0.75}$ | 0 | 2.39 (0.04) | 2.16 (0.02) | -8.60 (1.57) |
| | -0.3 | 2.37 (0.05) | 2.23 (0.03) | -3.60 (2.39) |
| | -0.6 | 2.44 (0.05) | 2.22 (0.03) | -7.59 (2.04) |
| | 0.3 | 2.40 (0.05) | 2.18 (0.04) | -7.97 (2.08) |
| | 0.6 | 2.42 (0.04) | 2.14 (0.03) | -10.66 (1.73) |
| $H_{0:1}$ | 0 | 2.35 (0.05) | 2.27 (0.03) | -1.07 (2.31) |
| | -0.3 | 2.42 (0.04) | 2.22 (0.03) | -7.01 (1.93) |
| | -0.6 | 2.43 (0.05) | 2.17 (0.03) | -8.76 (2.01) |
| | 0.3 | 2.42 (0.05) | 2.19 (0.03) | -7.78 (1.98) |
| | 0.6 | 2.41 (0.05) | 2.12 (0.03) | -10.2 (2.06) |

*Inbreeding rate in 36 BQs scenarios and comparison to 24 BQs scenarios*

**Table S5: Inbreeding rate in 36 BQs scenarios in the Base and Alt breeding scheme and relative difference with 24 BQs scenarios, for a null between-effects genetic correlation.**

| **Breeding goal** | $r_{T1,T2}$ | **Inbreeding rate in 36 BQs scenarios (%)** | | **Relative difference between 36 and 24 BQs scenarios (%)** | | **Relative difference between 36 BQs scenarios in Alt and 24 BQs scenarios in Base (%)** |
|---|---|---|---|---|---|---|
| | | Base breeding scheme | Alt breeding scheme | Base breeding scheme | Alt breeding scheme | |
| **$H_{1:0}$** | 0 | 1.53 (0.03) | 1.48 (0.02) | -32.75 (2.01) | -28.61 (1.57) | -13.13 (2.33) |
| | -0.6 | 1.55 (0.03) | 1.37 (0.02) | -31.07 (1.86) | -32.93 (1.49) | -18.51 (1.97) |
| | 0.6 | 1.52 (0.03) | 1.45 (0.02) | -30.37 (2.26) | -27.64 (1.53) | -11.21 (2.55) |
| **$H_{0.75:0.25}$** | 0 | 1.51 (0.03) | 1.43 (0.02) | -34.16 (2.14) | -29.42 (1.57) | -16.88 (2.39) |
| | -0.6 | 1.57 (0.03) | 1.43 (0.02) | -31.87 (1.74) | -30.61 (1.46) | -17.50 (1.75) |
| | 0.6 | 1.52 (0.03) | 1.42 (0.02) | -32.07 (1.82) | -28.33 (1.39) | -15.27 (1.91) |
| **$H_{0.5:0.5}$** | 0 | 1.60 (0.03) | 1.49 (0.03) | -32.78 (2.04) | -29.46 (1.57) | -16.68 (2.4) |
| | -0.6 | 1.66 (0.03) | 1.49 (0.02) | -31.78 (1.97) | -30.16 (1.42) | -18.12 (2.14) |
| | 0.6 | 1.52 (0.03) | 1.40 (0.02) | -31.98 (2.10) | -31.40 (1.63) | -16.37 (2.44) |
| **$H_{0.25:0.75}$** | 0 | 1.58 (0.04) | 1.64 (0.03) | -33.34 (2.01) | -24.09 (1.73) | -7.82 (2.42) |
| | -0.6 | 1.59 (0.03) | 1.58 (0.03) | -31.23 (2.04) | -28.08 (1.62) | -9.13 (2.62) |
| | 0.6 | 1.49 (0.04) | 1.46 (0.03) | -34.99 (2.10) | -28.93 (1.76) | -15.57 (2.47) |
| **$H_{0:1}$** | 0 | 1.60 (0.03) | 1.65 (0.02) | -30.99 (1.72) | -26.75 (1.59) | -5.04 (2.21) |
| | -0.6 | 1.56 (0.03) | 1.46 (0.02) | -32.02 (2.01) | -30.88 (1.49) | -14.85 (2.26) |
| | 0.6 | 1.52 (0.03) | 1.47 (0.02) | -34.04 (1.83) | -30.78 (1.49) | -14.95 (2.13) |

Standard errors (SE) are shown in brackets. SE for relative differences ($\mathrm{SE_R}$) across 24 and 36 BQs simulations were estimated using propagation of uncertainties from SE of means (μ) obtained within 24 ($\mathrm{SE_{24}}$) and within 36 BQs ($\mathrm{SE_{36}}$) simulations:

$$\mathrm{SE_R} = \frac{1}{\mu_{24}}\sqrt{\mathrm{SE}_{36}^2 + \left(\frac{\mu_{36}\cdot \mathrm{SE}_{24}}{\mu_{24}}\right)^2}$$

**Table S6: Inbreeding rate in 36 BQs scenarios in the Base and Alt breeding scheme and relative difference with 24 BQs scenarios, for a between-effects genetic correlation of -0.5.**

| Breeding goal | $r_{T1,T2}$ | Inbreeding rate in 36 BQs scenarios (%) | | Relative difference between 36 and 24 BQs scenarios (%) | | Relative difference between 36 BQs scenarios in Alt and 24 BQs scenarios in Base (%) |
|---|---|---|---|---|---|---|
| | | Base breeding scheme | Alt breeding scheme | Base breeding scheme | Alt breeding scheme | |
| $H_{1:0}$ | 0 | 1.64 (0.04) | 1.53 (0.03) | -32.61 (2.01) | -30.10 (1.65) | -16.28 (2.20) |
| | -0.6 | 1.69 (0.03) | 1.54 (0.03) | -28.49 (1.91) | -27.48 (1.54) | -13.24 (2.14) |
| | 0.6 | 1.66 (0.03) | 1.50 (0.02) | -29.65 (1.83) | -28.80 (1.50) | -15.15 (2.02) |
| $H_{0.75:0.25}$ | 0 | 1.74 (0.04) | 1.55 (0.03) | -27.02 (2.12) | -26.66 (1.67) | -12.87 (2.33) |
| | -0.6 | 1.58 (0.04) | 1.50 (0.03) | -32.53 (2.13) | -31.18 (1.48) | -14.06 (2.32) |
| | 0.6 | 1.65 (0.03) | 1.52 (0.03) | -29.46 (2.03) | -29.17 (1.64) | -13.37 (2.29) |
| $H_{0.5:0.5}$ | 0 | 1.62 (0.04) | 1.55 (0.02) | -33.22 (2.08) | -27.88 (1.50) | -15.21 (2.27) |
| | -0.6 | 1.69 (0.03) | 1.62 (0.03) | -32.82 (1.69) | -26.60 (1.69) | -14.26 (2.27) |
| | 0.6 | 1.58 (0.04) | 1.51 (0.03) | -33.86 (2.02) | -29.61 (1.69) | -15.33 (2.07) |
| $H_{0.25:0.75}$ | 0 | 1.63 (0.04) | 1.57 (0.03) | -31.82 (1.94) | -27.28 (1.53) | -12.57 (2.21) |
| | -0.6 | 1.68 (0.03) | 1.64 (0.03) | -31.09 (1.97) | -26.26 (1.61) | -10.65 (2.27) |
| | 0.6 | 1.62 (0.03) | 1.53 (0.02) | -33.30 (1.74) | -28.52 (1.45) | -15.67 (1.70) |
| $H_{0:1}$ | 0 | 1.63 (0.03) | 1.66 (0.03) | -30.48 (2.03) | -26.84 (1.55) | -5.45 (2.48) |
| | -0.6 | 1.69 (0.04) | 1.54 (0.03) | -30.30 (2.18) | -28.89 (1.50) | -15.10 (2.25) |
| | 0.6 | 1.61 (0.03) | 1.49 (0.03) | -33.11 (2.06) | -29.94 (1.63) | -17.71 (2.32) |

Standard errors (SE) are shown in brackets. SE for relative differences ($SE_R$) across 24 and 36 BQs simulations were estimated using propagation of uncertainties from SE of means (μ) obtained within 24 ($SE_{24}$) and within 36 BQs ($SE_{36}$) simulations:

$$SE_R = \frac{1}{\mu_{24}} \sqrt{SE_{36}^2 + \left(\frac{\mu_{36} \cdot SE_{24}}{\mu_{24}}\right)^2}$$

*Genetic gain in 36 BQs scenarios and comparison to 24 BQs scenarios*

**Table S7: Increase in genetic gain for the breeding goal in 36 BQs scenarios in the Base and Alt breeding scheme and relative difference with 24 BQs scenarios, for a null between-effects genetic correlation.**

| Breeding goal | $r_{T1,T2}$ | Genetic gain in 36 BQs scenarios (%) | | Relative difference between 36 and 24 BQs scenarios (%) | | Relative difference between 36 BQs scenarios in Alt and 24 BQs scenarios in Base (%) |
|---|---|---|---|---|---|---|
| | | Base breeding scheme | Alt breeding scheme | Base breeding scheme | Alt breeding scheme | |
| $H_{1:0}$ | 0 | 31.57 (0.29) | 49.31 (0.30) | -9.58 (1.31) | -4.35 (0.96) | 41.25 (1.8) |
| | -0.6 | 32.85 (0.30) | 49.56 (0.30) | -6.46 (1.03) | -5.15 (0.95) | 41.13 (1.7) |
| | 0.6 | 33.04 (0.28) | 50.14 (0.36) | -6.79 (1.13) | -4.57 (1.07) | 41.44 (1.6) |
| $H_{0.75:0.25}$ | 0 | 25.30 (0.20) | 37.65 (0.29) | -6.82 (1.11) | -5.43 (0.94) | 38.68 (1.65) |
| | -0.6 | 19.62 (0.21) | 30.18 (0.18) | -7.92 (1.43) | -3.29 (1.03) | 41.59 (1.82) |
| | 0.6 | 31.16 (0.23) | 46.17 (0.18) | -7.17 (1.06) | -4.82 (0.83) | 37.54 (1.32) |
| $H_{0.5:0.5}$ | 0 | 22.83 (0.18) | 30.70 (0.23) | -8.77 (1.12) | -3.75 (1.10) | 22.65 (1.48) |
| | -0.6 | 12.90 (0.14) | 15.44 (0.18) | -5.48 (1.84) | -3.73 (1.93) | 13.17 (2.29) |
| | 0.6 | 30.32 (0.20) | 42.86 (0.29) | -6.82 (0.98) | -4.99 (1.03) | 31.70 (1.40) |
| $H_{0.25:0.75}$ | 0 | 25.63 (0.22) | 28.95 (0.32) | -5.73 (1.28) | 0.57 (1.80) | 6.46 (1.64) |
| | -0.6 | 20.48 (0.21) | 22.52 (0.29) | -4.45 (1.38) | 3.25 (1.88) | 5.09 (1.72) |
| | 0.6 | 30.98 (0.25) | 40.81 (0.33) | -8.36 (1.10) | -4.43 (1.16) | 20.72 (1.46) |
| $H_{0:1}$ | 0 | 32.19 (0.27) | 34.11 (0.48) | -7.75 (1.19) | 3.30 (2.23) | -2.24 (1.68) |
| | -0.6 | 33.30 (0.29) | 40.86 (0.33) | -7.08 (1.17) | -4.55 (1.34) | 13.99 (1.39) |
| | 0.6 | 33.31 (0.28) | 40.63 (0.41) | -6.10 (1.38) | -4.67 (1.27) | 14.56 (1.81) |

Standard errors (SE) are shown in brackets. SE for relative differences ($\mathrm{SE_R}$) across 24 and 36 BQs simulations were estimated using propagation of uncertainties from SE of means (μ) obtained within 24 ($\mathrm{SE_{24}}$) and within 36 BQs ($\mathrm{SE_{36}}$) simulations:

$$\mathrm{SE_R} = \frac{1}{\mu_{24}}\sqrt{\mathrm{SE}_{36}^2 + \left(\frac{\mu_{36}\cdot \mathrm{SE}_{24}}{\mu_{24}}\right)^2}$$

**Table S8: Increase in genetic gain for the breeding goal in 36 BQs scenarios in the Base and Alt breeding scheme and relative difference with 24 BQs scenarios, for a between-effects genetic correlation of -0.5.**

| Breeding goal | $r_{T1,T2}$ | Genetic gain in 36 BQs scenarios (%) | | Relative difference between 36 and 24 BQs scenarios (%) | | Relative difference between 36 BQs scenarios in Alt and 24 BQs scenarios in Base (%) |
|---|---|---|---|---|---|---|
| | | Base breeding scheme | Alt breeding scheme | Base breeding scheme | Alt breeding scheme | |
| **$H_{1:0}$** | 0 | 20.83 (0.25) | 31.8 (0.35) | -7.05 (1.66) | -3.38 (1.38) | 41.9 (2.44) |
| | -0.6 | 21.61 (0.26) | 32.46 (0.30) | -4.87 (1.58) | -2.95 (1.46) | 42.86 (2.11) |
| | 0.6 | 22.04 (0.23) | 32.22 (0.26) | -3.81 (1.62) | -3.34 (1.15) | 40.62 (2.18) |
| **$H_{0.75:0.25}$** | 0 | 16.54 (0.20) | 24.57 (0.19) | -6.63 (1.68) | -5.51 (1.12) | 38.7 (2.14) |
| | -0.6 | 12.76 (0.15) | 19.53 (0.18) | -4.08 (1.88) | -2.08 (1.34) | 46.82 (2.7) |
| | 0.6 | 20.65 (0.19) | 30.04 (0.23) | -3.56 (1.48) | -1.49 (1.28) | 40.33 (2.03) |
| **$H_{0.5:0.5}$** | 0 | 14.72 (0.17) | 18.92 (0.27) | -7.12 (1.55) | -2.47 (1.91) | 19.36 (2.23) |
| | -0.6 | 7.95 (0.14) | 9.15 (0.17) | -9.1 (2.27) | -0.94 (2.55) | 4.54 (2.69) |
| | 0.6 | 19.67 (0.18) | 27.23 (0.25) | -8.06 (1.41) | -3.16 (1.43) | 27.3 (1.95) |
| **$H_{0.25:0.75}$** | 0 | 16.98 (0.19) | 17.08 (0.32) | -3.89 (1.72) | 3.29 (2.70) | -3.3 (2.26) |
| | -0.6 | 12.68 (0.16) | 12.95 (0.22) | -4.45 (1.80) | 5.10 (2.70) | -2.47 (2.17) |
| | 0.6 | 20.6 (0.23) | 26.04 (0.25) | -4.99 (1.43) | -0.14 (1.64) | 20.07 (1.64) |
| **$H_{0:1}$** | 0 | 21.38 (0.25) | 19.69 (0.40) | -3.49 (1.71) | 5.92 (3.07) | -11.09 (2.16) |
| | -0.6 | 21.88 (0.25) | 25.04 (0.30) | -4.22 (1.56) | -2.85 (1.87) | 9.59 (1.83) |
| | 0.6 | 21.62 (0.22) | 25.08 (0.30) | -6.31 (1.51) | -2.42 (1.81) | 8.7 (1.88) |

Standard errors (SE) are shown in brackets. SE for relative differences ($\mathrm{SE_R}$) across 24 and 36 BQs simulations were estimated using propagation of uncertainties from SE of means (μ) obtained within 24 ($\mathrm{SE_{24}}$) and within 36 BQs ($\mathrm{SE_{36}}$) simulations:

$$\mathrm{SE_R} = \frac{1}{\mu_{24}}\sqrt{\mathrm{SE}_{36}^2 + \left(\frac{\mu_{36}\cdot \mathrm{SE}_{24}}{\mu_{24}}\right)^2}$$
